# Dachsous-Fat Signaling Shapes the Drosophila Wing through Mechanical Forces

**DOI:** 10.1101/2025.08.08.669334

**Authors:** Bipin Kumar Tripathi, Zhenru Zhou, Kenneth D. Irvine

**Author notes:** Department of Proteomic and Genomic Technologies, Genentech Inc, South San Francisco, California, USA.

## Abstract

Proper organ shape is critical for function. The *Drosophila* wing normally adopts an elongated shape, but mutations in the Dachsous-Fat pathway result in rounder wings. The mechanism by which this occurs has remained unclear. Here, we show that Ds-Fat signaling shapes the wing during the larval stage, rather than during pupal development when morphogenetic rearrangements transform the developing wing disc into the adult wing. We further find that Ds-Fat alters tissue wide stresses in the wing disc, and genetic manipulations that reduce cytoskeletal tension result in rounder wings, whereas increasing cytoskeletal tension produces more elongated wings. Reduced tension is also associated with less oriented growth during development. Notably, increased cytoskeletal tension partially rescues the rounder shape caused by *ds* knockdown. These results reveal a previously unrecognized mechanism by which Ds-Fat signaling determines wing shape, involving regulation of tissue tension to orient growth and shape the wing primordia during larval development.

## Introduction

Forming organs of the correct size and shape is crucial for their normal function, and congenital disorders in which organs do not form correctly have a substantial impact on human health. In many cases genes that are associated with congenital malformations play conserved roles in animal development reviewed in ^1,2^. Genes in the Dachsous-Fat pathway were first identified for their effects on the size and shape of wings and legs in *Drosophila* ^3–8^. Mutations in the human homologs of *Drosophila* Fat and Dachsous (Ds), FAT4 and DCHS1, can result in congenital diseases associated with organ malformation, including Van Maldergem syndrome, Hennekam syndrome, and Mitral Valve Prolapse ^9–11^. Gene targeted mutations in murine Fat4 or Dchs1 genes similarly result in defects in formation of multiple organs ^12–16^.

Dachsous and Fat are large cadherin family proteins that initiate intercellular signaling to control organ growth, shape and planar cell polarity (PCP) ^17–19^. Ds and Fat bind each other through their extracellular domains, with this binding modulated by Four-jointed (Fj)-mediated phosphorylation of cadherin domains ^20–23^. Ds, Fj, and in some cases, Fat, are expressed in gradients across tissues, and their differential expression and binding interactions leads to polarized membrane localization of Ds and Fat ^4,20,24–30^. Polarization of Ds and Fat leads to polarized membrane localization of a key downstream effector, Dachs ^5,28,29,31^. Dachs is an unconventional myosin protein that mediates connections of Ds-Fat to regulation of Hippo signaling and PCP ^5,27,28,32–36^.

Ds-Fat signaling plays key roles in many different organs but has been most intensively studied in the developing *Drosophila* wing. The wing develops from the wing imaginal disc, a cluster of ∼30 cells specified during embryogenesis that undergo extensive proliferation, growing ∼1000 fold during larval development ^37^. During pupal development, the wing disc undergoes morphogenetic changes including eversion, flattening, and expansion of the future wing blade to form the adult wing, wing hinge, and notum. The wing normally has an elongated, elliptical shape. This shape reflects both how the wing primordia, referred to as the wing pouch, grows within the developing larval wing disc, as well as morphogenetic processes that occur during pupal development. One key event within the pupa is a contraction of the wing hinge, which generates an anisotropic stress that pulls on the future wing blade and contributes to its elongation ^38,39^. Loss of *ds*, *fat*, or other core components of Ds-Fat signaling results in wings that are rounder than wild-type wings. A potential explanation for this was suggested by examination of growth and spindle orientation during larval development. Growth orientation can be visualized by making marked clones of cells, which tends to be oriented along the proximal-distal axis of the developing wing; this orientation correlates with a preferential orientation of mitotic spindles along the proximal-distal axis. This orientation of mitotic spindles, and of clone growth, is lost in *ds*, *fat* or *dachs* mutants, which led to the suggestion that Ds-Fat signaling controls growth orientation, and ultimately wing shape, through effects on spindle orientation ^40,41^.

The suggestion that Ds-Fat signaling control wing shape through effects on spindle orientation was called into question by the discovery that spindle orientations are randomized by mutation of *mud*, but clone growth and wing shape are nonetheless normal in *mud* mutants ^42^. How then might the effect of Ds-Fat on wing shape be explained? One clue comes from investigations of cellular contributions to tissue shear. As the wing disc grows during mid-third instar, three cellular behaviors account for most shear along the proximal distal axis: oriented cell divisions, oriented cell rearrangements, and oriented cell shape changes. Intriguingly, cellular analysis of live discs cultured ex vivo revealed that the relative contributions of these processes can vary between individual discs ^43^. Moreover, in *mud* mutant discs, the loss of oriented cell divisions was at least partially compensated for by an increased contribution of cell rearrangements to tissue shear ^42^. These observations suggest that the cellular behaviors observed could be interchangeable responses to tissue stress.

Here, we revisit the question of how Ds-Fat signaling influences wing shape. We confirm that Ds-Fat signaling acts during larval stages to control adult wing shape. Our investigations reveal that *ds* and *ft* mutants alter the shape of the wing pouch from the earliest stages of its formation. This initial alteration of wing pouch shape is shared by mutations that disrupt Hippo signaling, but alterations in Hippo signaling, or in canonical PCP, cannot explain the influence of Ds-Fat on wing shape. Instead, we find that loss of Ds or Fat alters tissue stresses within the wing pouch, and that these alterations are associated with changes in the distribution of non-muscle myosin II. We further show that direct alterations of cytoskeletal tension can alter wing shape, and can modulate the consequences of *ds* mutations on wing shape. Altogether, our observations support a hypothesis in which Ds-Fat signaling controls wing shape by patterning tissue stresses that shape the orientation of growth throughout larval development.

## RESULTS

### Dachsous and Fat are required during larval development for wing shape

Mutations in several components of the Ds-Fat pathway, including *ds*, result in rounder wings ^4,5,30,40,41,44^ (Figures 1A, B). This can be quantified by comparing the length of the wing to its width. To simplify these measurements, we fit a tracing of the outline of the wing to an ellipse and plotted the ratio of the major axis (length) of the ellipse to its minor axis (width) (Figure 1C). Using this approach, we measured a major/minor axis ratio of 2.1 for wild-type wings, and 1.6 for *ds* mutant wings (Figure. 1D). Ds and Fat influence growth orientation during larval development ^40,41^, but also influence oriented cell behaviors during pupal development ^45–47^. To distinguish larval versus pupal contributions of *ds* and *fat* to adult wing shape, we used conditional expression of RNAi transgenes. *UAS-RNAi*-*fat* or *UAS-RNAi*-*ds* transgenes were expressed under *nub-Gal4* control, which begins to be expressed in the developing wing blade and distal hinge during the second larval instar ^48^. Temporal control of expression was provided using a ubiquitously-expressed temperature-sensitive GAL80 (tub-GAL80^ts^) to antagonize GAL4 ^49^. When Fat or Ds were knocked down throughout most of larval development by keeping flies continuously at 29°C (inactivating Gal80^ts^), then rounder adult wings were generated, similar to those observed in *ds* mutants (Figure 1E-G, P). When *ds* or *fat* RNAi was suppressed by keeping flies continuously at 18°C, then wing shapes similar to those in wild-type controls were observed (Figures 1H, I, P). Knockdown of Fat or Ds from embryonic through late 3^rd^ instar or early pupal stages (Figure 1J-M, P) resulted in rounder wings, like in *ds* mutants. Conversely, knockdown of Fat or Ds from early pupal to adult stages resulted in wing shapes similar to wild-type controls (Figure 1N-P). Together, these experiments indicate that Ds and Ft are required during larval development, but not during pupal development, for normal wing shape.

**Figure 1:**
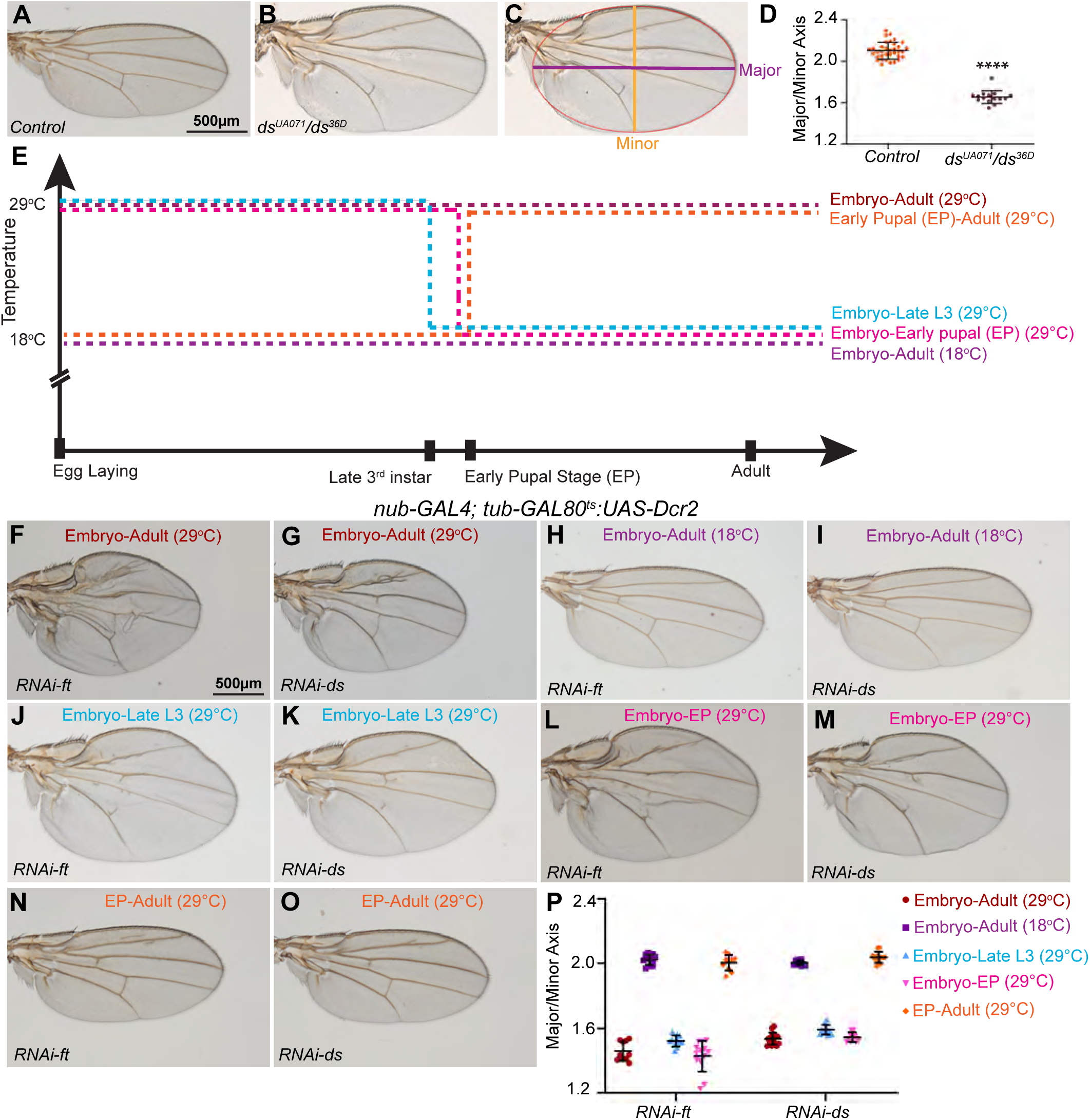
Effect of temporal knockdown of *ft* and *ds* on wing shape. **(A-C)** Male wings from *w*^1118^ (A), *ds^UA^*^071^*/ds^36D^* (B,C). In C the fitted ellipse (red) used to measure the major axis (purple line) and minor axis (orange line) is overlayed. Scale bar=500 µm. **(D)** Histogram illustrating the Major/Minor axis ratio of control (*w*^1118^, n=35) and *ds^UA^*^071^*/ds^36D^* (n=15). Error bar indicates mean±s.d., the significance of differences by t-test is indicated by black asterisks. **(E)** Schematic for temperature shift experiment to knockdown Ft and Ds using the *GAL4/GAL80^ts^* system at different developmental stages. **(F-O)** Knockdown of Ft from embryo to adult (F; n=10), Ds from embryo to adult (G; n=14), Ft control (H; n=11), Ds control (I; n=10), Ft from embryo to late 3^rd^ instar (J; n=11), Ds from embryo to late 3^rd^ instar (K; n=11), Ft from embryo to early pupal (L; n=13), Ds from embryo to early pupal (M; n=11), Ft from early pupal to adult (N; n=10), Ds from early pupal to adult (O; n=12). Scale bar=500 µm. **(P)** Histogram quantifying wing shapes for wings as shown in F-O. Error bar indicates mean±s.d.

### Ds-Fat signaling regulates the shape of the wing pouch in larval wing discs

The adult wing blade develops from a central region of the ventral half of the larval wing imaginal disc referred to as the wing pouch, which is demarcated by folds that form in the future wing hinge^37^. During metamorphosis, the wing pouch everts and folds in half, such that the dorsal-ventral midline of the wing pouch becomes the margin (edge) of the adult wing (Figure 2A). To investigate whether the altered shape of the adult wing is reflected in the shape of the wing pouch, we compared wing pouch shape in wild-type versus *ds* or *fat* mutant wing discs. We used expression of Wingless (Wg) to analyze wing pouch shape, as it is expressed in two rings of cells in the developing hinge as well as in a line of cells along the D-V compartment boundary (Figure 2B). We compared the D-V/A-P ratio, with the D-V length defined as the length of Wg expression along the D-V boundary within the inner ring of Wg expression (Figure 2A), and the A-P length defined as the width of the inner ring at its center (roughly along the A-P compartment boundary) from top (dorsal) to bottom (ventral). In wild-type wing discs at late third instar, the D-V/A-P ratio is ∼1.7 (Figure 2B,E). In contrast, in *ds* mutant wing discs, the D-V/A-P ratio is ∼2.4 and in *ft* mutant wing discs the D-V/A-P ratio is ∼2.5 (Figure 2C-E). As the D-V length in the wing disc corresponds to the circumference of the adult wing, while the A-P length corresponds to the proximal-distal length of the adult wing, a larger D-V/A-P ratio in the wing disc corresponds to a wider adult wing.

**Figure 2:**
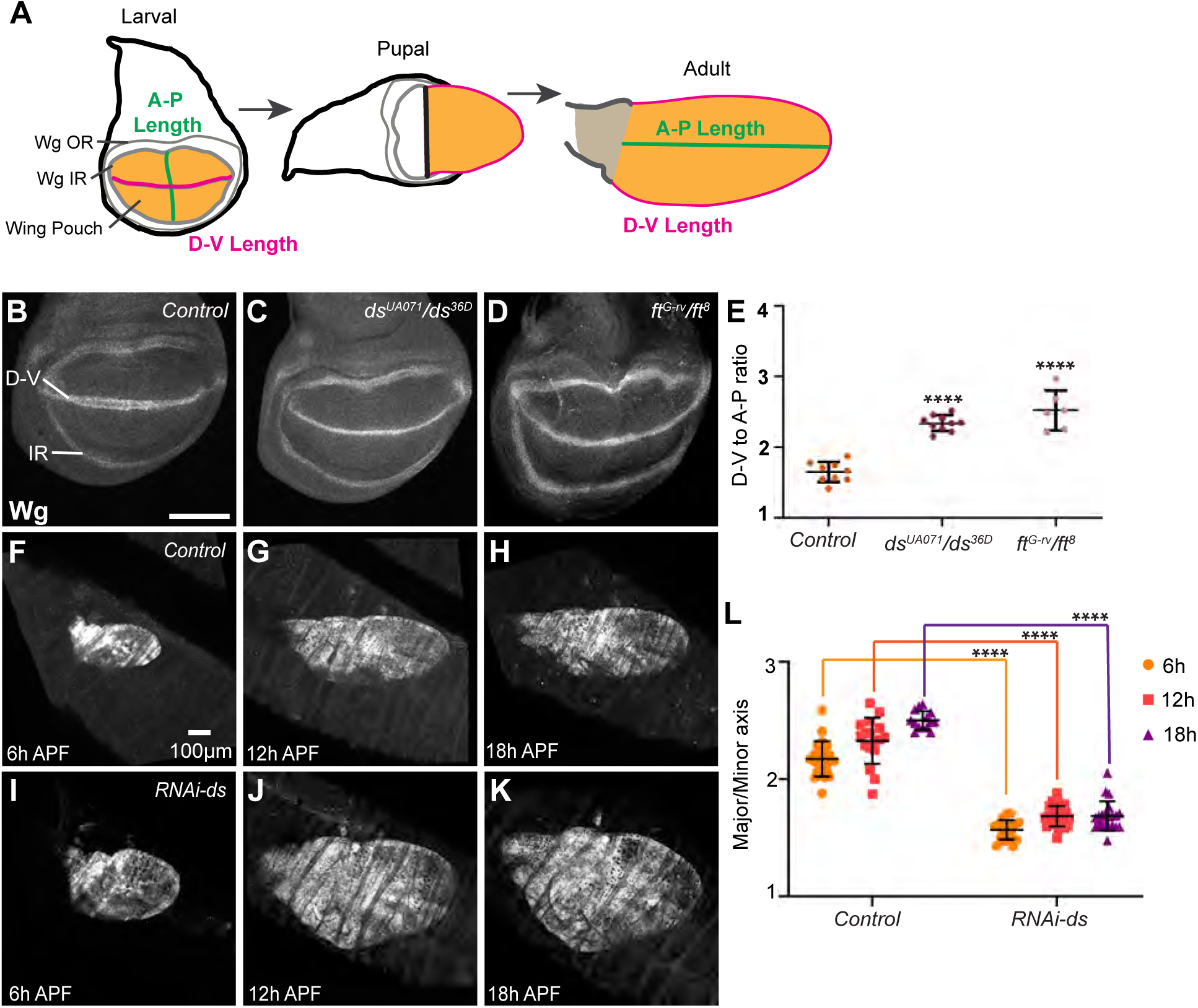
Altered wing pouch shape in *ds* and *ft* mutants. (A) Schematic showing relationship between wing disc and adult wing, with approximate locations of wing pouch (orange), D-V length (magenta) and A-P length (green), Wg outer (OR) and inner (IR) rings indicated. **(B-D)** Late 3^rd^ instar wing discs stained for Wg from *w*^1118^ (B; control, n=9), *ds^UA^*^071^*/ds^36D^* (C; n=10), and *ft^G-rv^/ft*^8^ (D; n=6). Scale bar=100 µm. **(E)** Histogram illustrating the wing pouch shape for wing discs of genotypes shown in B-D. Error bar indicates mean±s.d., the significance of differences relative to *w*^1118^, calculated using one-way ANOVA on measurements from the number of wing discs indicated above is indicated by black asterisks. **(F-K)** Pupal wings from *nub-GAL4:UAS-Dcr2/+; UAS-mCD8:RFP/+* (Control, F-H) or *nub-GAL4:UAS-Dcr2/RNAi-ds; UAS-mCD8:RFP/+* (I-K) at 6h (F, n=23, I, n=19), 12h (G, n=17, J, n=25), and 18h (H, n=14, K, n=22) APF. Scale bar=100 µm. **(L)** Histograms showing measurements of pupal wing shape. Error bar indicates mean±s.d., the significance of differences at the same developmental stage is indicated by black asterisks, calculated by t-test on measurements from the number of wings indicated above.

### Influence of Ds-Fat signaling on pupal wing shape

To further examine the conclusion that altered larval wing pouch shape accounts for altered adult wing shape, we examined developing pupal wings from wild-type and *ds* RNAi knockdown animals. For these experiments, we used nub-GAL4 driving the expression of UAS-mCD8:RFP to label developing pupal wings. To minimize disturbance of pupal wing morphology, developing wings were imaged live directly through the pupal case. We found that at the beginning of pupal development (white prepupae) the three-dimensional shape of the everting wing made it difficult to reliably measure wing shape. However, by 6 hours (h) after puparium formation (APF), the wing has everted and folded into opposing dorsal and ventral surfaces, and even at 6h APF, *ds* RNAi wings were substantially wider than control wings (Figure 2F,I,L). The difference in shape was maintained as the wings flatten and expand (Figure 2G,H,J-L). These observations are consistent with the conclusion that *ds* acts during larval stages to regulate wing shape.

### Influence of Ds-Fat signaling on wing pouch shape is visible at early third instar

Ds-Fat signaling can influence the organization of growth during larval development, as the shapes of clones growing in *ds*, *fat* or *dachs* mutant wings are more rounded than clones growing in wild-type wings ^40,41^. If the altered shape of the larval wing disc were solely a consequence of altered patterns of growth, then we reasoned that this altered shape would arise gradually over third instar, in conjunction with disc growth. To investigate this possibility, we examined the developmental profile of wing pouch shape throughout the third larval instar (at 72, 84, 96, 108, and 120 h after egg laying [AEL]), using analysis of the D-V/A-P ratio in wing discs stained for Wg expression. In wild-type wing discs, the D-V/A-P ratio was relatively consistent from the earliest time that a ring of Wg expression in the hinge could be clearly identified (1.8 early third instar, 72 h AEL) through the end of the third larval instar (1.7 at 120 h AEL), and this slight decrease was not statistically significant (Figure 3A, D). Contrary to our expectations, in *ds* or *ft* mutants the D-V/A-P ratio was already abnormal at 72 h AEL, at 2.6 for *ds* and 2.9 for *ft* (Figure 3 B-D). The D-V/A-P ratio in *ds* or *fat* mutants reduced slightly as the discs grew but remained significantly higher than that in wild-type controls throughout third instar. These observations indicate that Ds-Fat signaling influences wing shape during the initial formation of the wing pouch, in addition to its effects during wing growth.

**Figure 3:**
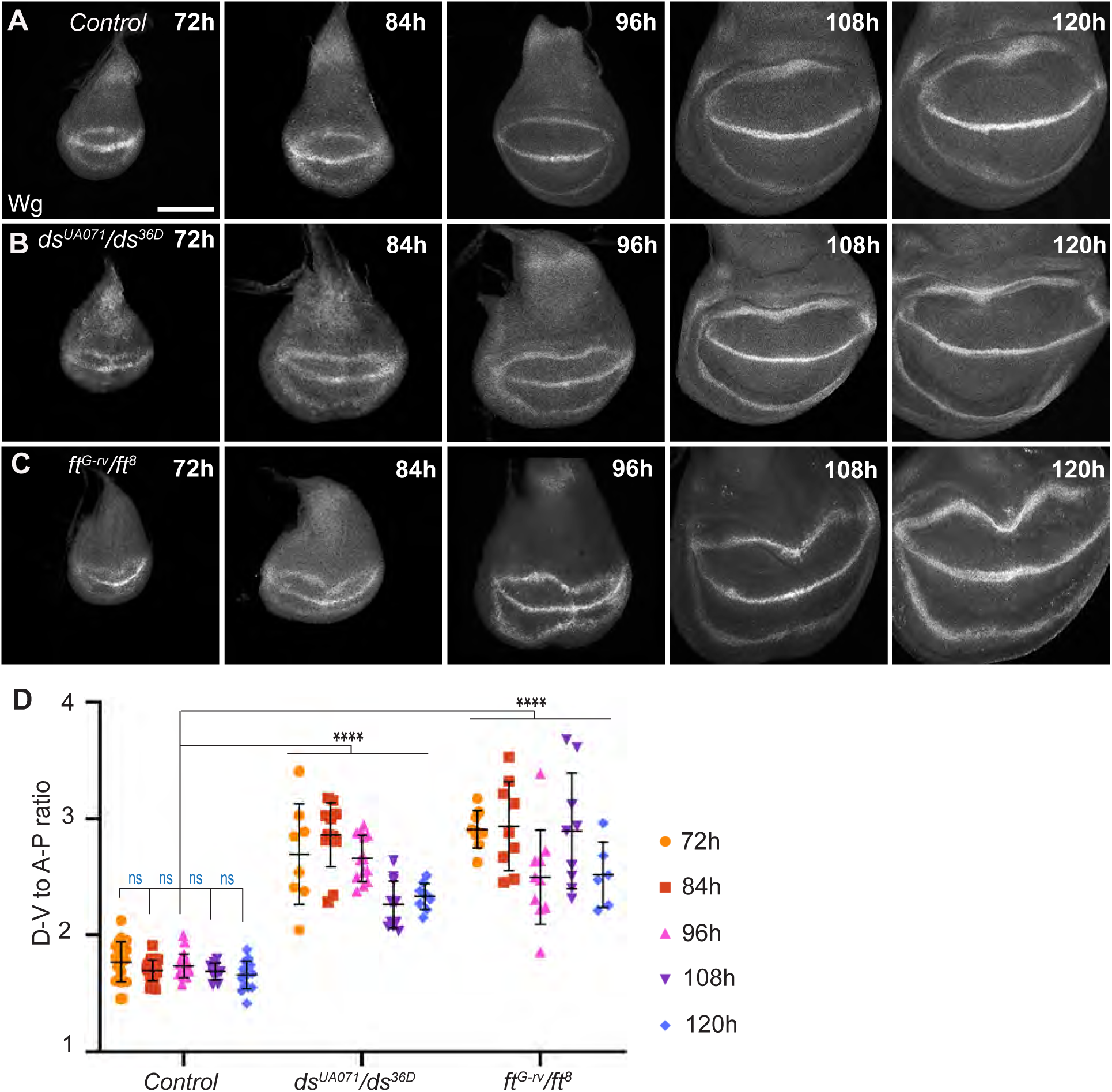
*ds* and *ft* mutants have altered wing pouch shapes throughout third instar. **(A-C)** Wing discs stained for Wg from **(A)** *w*^1118^**, (B)** *ds^UA^*^071^*/ds^36D^***, (C)** *ft^G-rv^/ft*^8^ at 72h (n=23 *w,* 8 *ds*, 9 *ft*), 84h (n=22 *w*, 13 *ds*, 9 *ft*), 96h (n=20 *w*, 13 *ds*, 10 *ft*), 108h (n=10 *w*, 11 *ds*, 9 *ft*), and 120h (n=15 *w*, 9 *ds*, 6 *ft*) AEL. Scale bar=100 µm. **(D)** Graph of wing pouch shape for discs described in A-C. Error bar indicates mean±s.d., and significance of differences for different developmental stages of *w*^1118^ is relative to its 72h stage and indicated as ns in blue; the significance of differences relative to *w*^1118^ at same developmental stage is indicated by black asterisks and were calculated using one-way ANOVA on measurements from the number of samples indicated above.

### The influence of Ds-Fat on wing shape is not explained by Hippo signaling

Ds-Fat signaling regulates distinct downstream processes: Hippo signaling and PCP. To investigate whether effects on Hippo signaling contribute to the influence of Ds-Fat on wing and wing disc shape, we examined flies mutant for the Hippo pathway regulator *expanded (ex)*, as adult flies can be recovered from animals mutant for *ex^e^*^1^, an amorphic allele of *ex* ^50^. *ex^e^*^1^ adult wings are rounder than wild-type wings, with a major/minor axis ratio of 1.8 (Figure S1A,B,D), but not as round as *ds* or *fat* mutant or RNAi wings (Figure 1B,D). Null mutations of *hippo* (*hpo*) are lethal, but wings from animals in which *hpo* was knocked down during wing development by RNAi expression under nub-Gal4 control were similarly intermediate in shape between wild-type controls and *ds* or *fat* mutants or RNAi (Figure S1C, D), with a major/minor axis ratio of 1.9. These observations suggest that Hippo signaling contributes to, but does not fully explain, the influence of *ds* or *fat* on adult wing shape.

We also examined the consequences of impairing Hippo signaling on wing disc shape. In *ex^e^*^1^ mutants, or in animals mutant for *wts* (*wts^P^*^2^*/ wts^X^*^1^), a hypomorphic combination of *wts* alleles, we observed that the initial wing disc shape at early third instar is elongated along the D-V boundary (Figure S1E-G), as observed in *ds* or *fat* mutants (Figure 3B,C). This suggests that the initial elongation of the wing pouch in *ds* or *ft* mutants could be due to impairment of Hippo signaling. However, as the wing disc grows, the D-V/A-P ratio in *ex^e^*^1^ or *wts* mutants gradually declines, such that by the end of third instar it is similar to that in wild-type controls (Figure S1E-G). These results imply that Ds-Fat influences wing pouch shape during wing disc growth separately from its effects on Hippo signaling.

During pupal development, the hinge region of the developing wing contracts ^46^. As the distal region of the wing is attached to apical extracellular matrix (ecm), this contraction contributes to elongation of the wing blade ^38,39^. This led us to consider whether the increased size of the wing in *ex* mutants or *hpo* RNAi might be a factor in their rounder adult wing shape. To investigate this, we increased growth through a distinct mechanism, by expressing an activated form of the Insulin receptor (*InR^CA^*) under *nub-GAL4* control. Expression of *InR^CA^* in the wing under *nub-Gal4* control results in larger adult wings, similar in size to those generated by Ft knockdown (Figure S2A-D). However, instead of being rounder, adult wings expressing *InR^CA^* were actually slightly more elongated than control wings (Figure S2A,B, E). Wing discs expressing *InR^CA^* were similar to controls in shape during early larval development (72h-96h AEL), but as the wing disc grows the D-V/A-P ratio in *UAS-InR^CA^ nub-GAL4* declines slightly, such that by the end of third instar the wing pouch is elongated along the A-P axis compared to controls (Figure S2F, G), which correlates with the observed elongation of adult wings (Figure S2B, E).

### Over-expression of Ds or Fat also alters wing pouch shape

Over-expression of Fat or Ds, or mutation of the downstream effector *dachs*, is associated with formation of wings that are both smaller and rounder than wild-type wings (Figure S3A-C, E) ^5,20,31^. To investigate whether the altered adult wing shape is reflected in altered wing pouch shape in the larval disc, we used *nub-Gal4* to drive expression of *UAS-fat* during wing development. Over-expression of Fat was associated with a higher D-V/A-P ratio, 1.9 even at 72h AEL, and the D-V/A-P ratio increased further during wing disc growth (2.1 at 120 h AEL) (Figure S3F, H). Over-expression of Ft is associated with removal of Dachs from cell membranes ^5^, and we also observed an elevated D-V/A-P ratio in *dachs* mutant wing discs throughout larval development (Figures S3G, I). The altered shape of the wing pouch in these genotypes is consistent with the rounder adult wings that they form. Notably, other genotypes that reduce Hippo signaling, such as downregulation of *jub* or *zyx*, or over-expression of *wts*, are associated with narrower adult wings rather than wider adult wings (Figure S3D, E)^51^. This implies that the influence of *fat* or *dachs* on wing shape is not due to their impact on Hippo signaling.

### Dachsous regulates patterns of tissue stress

The determination that randomizing spindle orientation does not alter wing shape ^42^, together with observations that variations in the contribution of spindle orientation to shear during wing disc growth can be compensated for by cell rearrangements or cell shape changes ^42,43^, suggest that the primary influence of *ds* and *fat* might be on stress patterns in the developing wing disc. To investigate whether stress patterns are altered in *ds* mutants, we used laser cutting of cell junctions ^52,53^. We made circular cuts with a 10 µm radius (encompassing ∼ 25-30 cells) in wing discs at 108h AEL. If the cut region is under tension, then the outer edge will expand as it is pulled by neighboring cells, with the extent of expansion correlating with the level of external tension and the shape revealing whether this tension is isotropic (yielding circular expansion) or anisotropic (yielding elliptical expansion). We made cuts in proximal or D-V boundary regions (Figure 4A) and measured the size and shape of the cut regions at their maximal expansion (∼1 minute after cutting). In proximal regions of wild-type controls, cut expansion after circular ablation resulted in an enlarged elliptical shape, revealing anisotropic stress (Figure 4B, F, G). Near the D-V boundary of wild-type controls, cut expansion after circular ablation also resulted in an enlarged elliptical shape, of similar size but less elliptical than in proximal regions (Figures 4D,H,I and S4A,C). When we made cuts in proximal regions of *ds* mutant wing discs, cut expansion resulted in shapes that were smaller and rounder than those in wild type discs (Figures 4C,F,G and S4B,D). Thus, in this region of *ds* mutants, tissue stress is lower and more isotropic. Conversely, near the D-V boundary, cut expansion after circular ablation in *ds* mutants resulted in enlarged areas that are larger and more elliptical than those in wild type discs (Figures 4E,H,I and S4B, D). Thus, in this region of *ds* mutants, tissue stress is higher and more anisotropic. Stress anisotropy often correlates with cells shape, with anisotropic stress associated with more elongated cells. Consistent with this, quantification of cell eccentricity revealed that cells are less elongated in proximal regions in *ds* mutant wing discs as compared to wild-type controls (Figures 4B,C,J,K).

**Figure 4:**
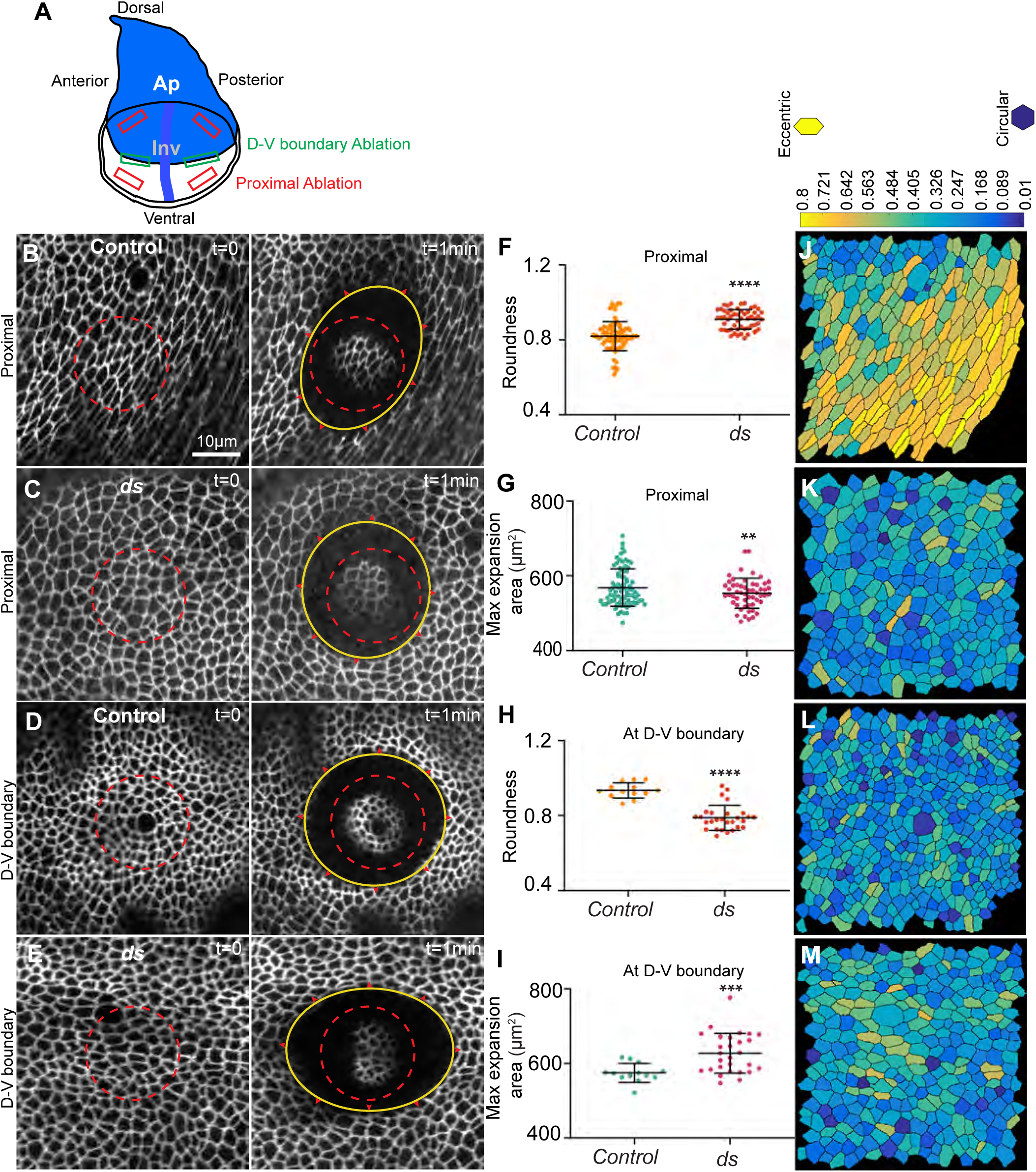
Mapping stress in *ds* mutants by circular ablation. **(A)** Schematic of late third instar wing disc, with expression patterns of Ap (blue) and Inv (dark blue), and approximate regions for circular ablation in the proximal wing pouch (red) and near the D-V boundary (green) indicated. **(B)** Wing disc expressing Ecad:GFP at 108h from control tissue (*Ecad:GFP/ inv-BFP ap-BFP*) showing the tissue before (t=0) and after (t=1min) a proximal circular ablation (n=83). **(C)** Wing disc expressing Ecad:GFP at 108h from *ds* mutant (*ds^UA^*^071^*/ds^36D^ ;Ecad:GFP/ inv-BFP ap-BFP*) showing the tissue before and after a proximal circular ablation (n=54). **(D)** Wing disc expressing Ecad:GFP at 108h from *control* showing the tissue before and after a circular ablation at D-V boundary (n=13). **(E)** Wing disc expressing Ecad:GFP at 108h from *ds* mutant showing the tissue before and after a circular ablation at D-V boundary (n=28). Scale bar=10 µm. The initial circular ablation region is marked by dotted red circle; the outer edge of the cut after 1 min is marked by a solid yellow ellipse. **(F-I)** Histograms plotting shape (F, H) and size (G, I) of the cut regions at their maximal expansion. Error bars indicate mean±s.d., the significance of differences relative to control tissue by t-tests on measurements from the number of samples indicated above is indicated by black asterisks. **(J-M)** Heatmaps showing cell eccentricity on segmented cells (from B-E), colored according to eccentricity using the scale at top.

### Influence of Ds-Fat signaling on myosin localization

The distribution of myosin is a key driver of, and generally correlates with, tension in the actin cytoskeleton. To determine whether the altered stress patterns we detected in *ds* mutants could be explained by changes in myosin distribution, we examined Sqh:GFP, a GFP-tagged myosin light chain (encoded in *Drosophila* by the *spaghetti squash* gene, *sqh*). Levels of Sqh:GFP at cell-cell junctions were quantified and compared with levels of E-cadherin (E-cad). Compared to control wing discs at 108h AEL, *ds* mutant discs have relatively higher junctional myosin near the D-V boundary, and lower junctional myosin in the rest of the wing pouch (Figure 5 A,B), consistent with our observations of higher tissue tension near the D-V boundary and lower tissue tension in proximal regions. To confirm these differences, we also examined myosin distribution in wing discs in which *ds* was knocked down in posterior cells under *hh-Gal4* control, leaving anterior cells as an internal control. This experiment also revealed increased levels of junctional myosin near the D-V boundary and reduced levels of junctional myosin in proximal regions in the *ds* knockdown compartment (Figure 5 C,D).

**Figure 5:**
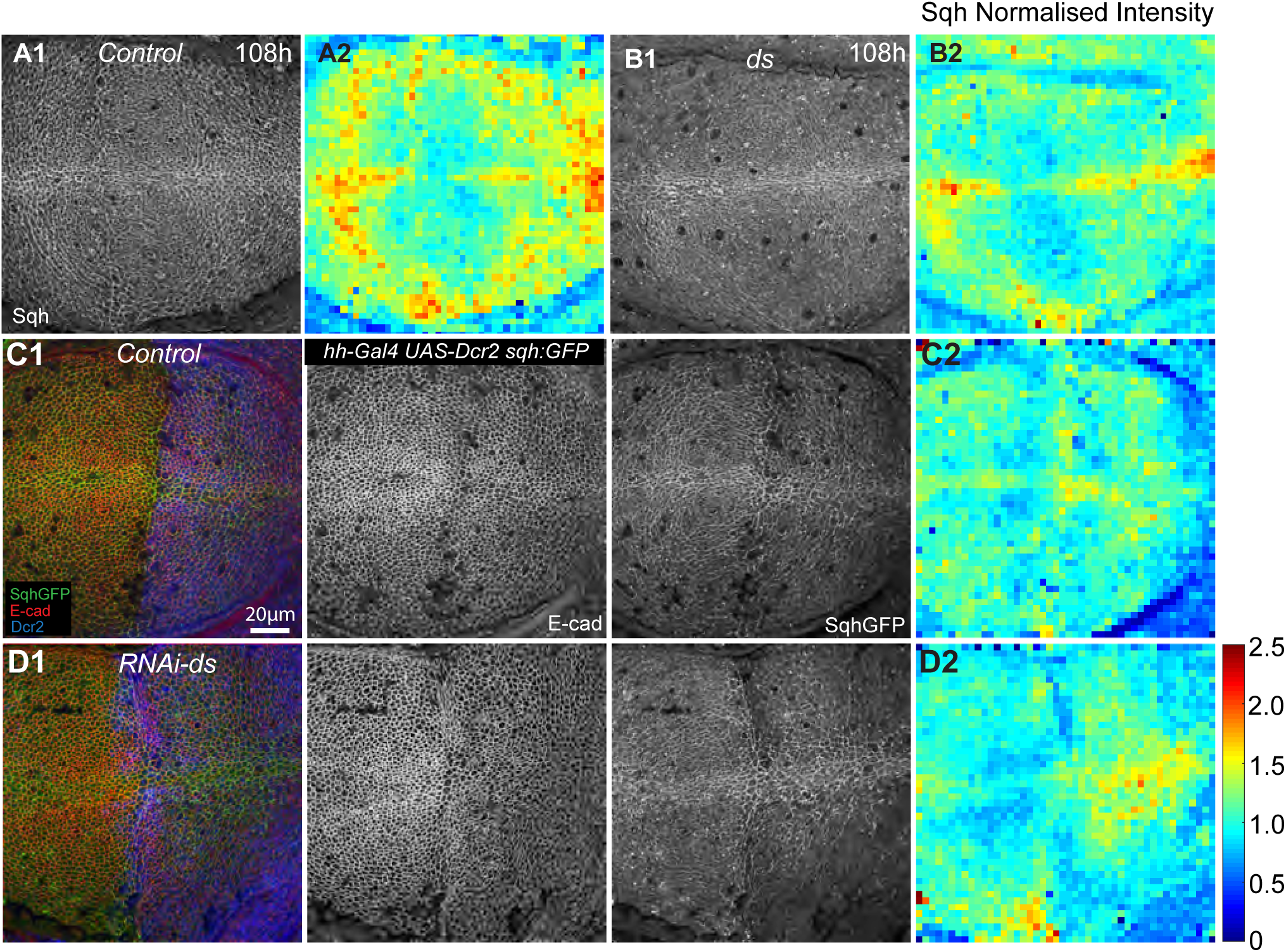
Altered myosin in *ds* mutants. Wing discs expressing Sqh:GFP at 108h from **(A1)** control (Sqh:GFP/+) and **(B1)** *ds^UA^*^071^*/ds^36D^; Sqh:GFP/+.* **(A2 and B2)** Heat map of Sqh:GFP intensity normalized to E-cad from A1 and B1 respectively. **(C1 and D1)** Wing discs expressing Sqh:GFP along with *hh-GAL4:UAS-Dcr2* in controls (C1), and in *RNAi-ds* (D1). Scale bar: 20 µm. **(C2 and D2)** Heat map of Sqh:GFP intensity normalized to E-cad from C1 and D1, respectively. Scale for the heat map is on the side, ranges from 0 to 2.5.

### Changes in cytoskeletal tension alter wing shape and growth orientation

Our examination of myosin distribution and tissue stress suggest that the Ds-Fat pathway alters wing shape by modulating tissue stress patterns. To further evaluate this hypothesis, we examined the consequences of genetically altering cytoskeletal tension on wing shape. We note that earlier studies suggest that altered tension can impact wing shape. For example, when Rho-associated protein kinase (Rok), which phosphorylates and activates myosin ^54^, is knocked down throughout the developing wing (UAS-RNAi-rok nub-Gal4) then wings appear smaller and rounder ^55^. We have extended these observations by measuring wing shapes and by analyzing the shape of the wing pouch during larval development. Quantitation confirmed that knockdown of *rok* results in rounder wings (Figure 6B, D). Conversely, increasing Rok activity by expressing an activated form of Rok comprising the catalytic domain (Rok.CA) resulted in flies with elongated wings (Figure 6C, D). To examine if these effects on adult wing shape are also reflected in larval wing pouch shape we measured the D-V/A-P ratio in wing discs from 72-120 h AEL. Decreasing Rok levels using RNAi-rok resulted in an increased D-V/A-P ratio, which was visible from early larval stages (72h AEL) and remained elevated throughout larval development (Figures 6F, H). Conversely, increasing Rok activity in the wing discs resulted in a slightly decreased D-V/A-P ratio compared to controls, and the difference was statistically significant at later stages of development (Figure 6 G, H).

**Figure 6:**
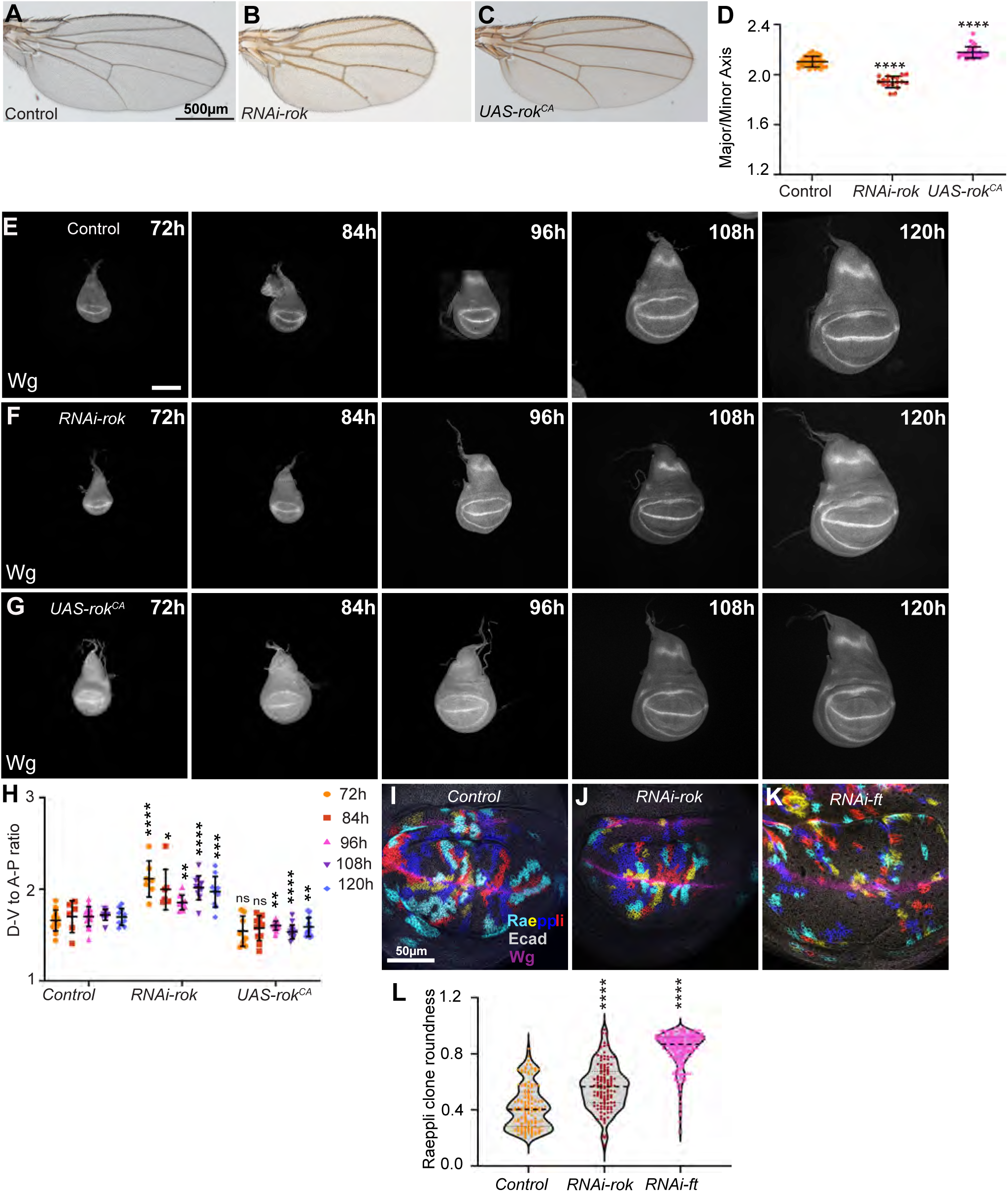
Influence of cytoskeletal tension on wing shape and growth orientation. **(A-C)** Male wings from *nub-GAL4:UAS-Dcr2/+* (A; n=32), *nub-GAL4:UAS-Dcr2/RNAi-rok* (B; n=18), and *nub-GAL4:UAS-Dcr2/UAS-rok^CA^* (C; n=33). Scale bar=500 µm. **(D)** Histogram quantifying adult wing shape for wings described in A-C. Error bar indicates mean±s.d., the significance of differences relative to control, calculated by one-way ANOVA, is indicated by asterisks. **(E-G)** Wing discs stained for Wg from **(E)** *nub-GAL4:UAS-Dcr2/+* at 72h AEL (n=14), 84h AEL (n=6), 96h AEL (n=15), 108h AEL (n=11), and 120h AEL (n=10), **(F)** *nub-GAL4:UAS-Dcr2/RNAi-rok* at 72h AEL (n=7), 84h AEL (n=7), 96h AEL (n=9), 108h AEL (n=11), and 120h AEL (n=10) **(G)** *nub-GAL4:UAS-Dcr2/UAS-rok^CA^* at 72h AEL (n=8), 84h AEL (n=11), 96h AEL (n=15), 108h AEL (n=14) and 120h AEL (n=18 Scale bar=100 µm. **(H)** Histogram quantifying wing pouch shape for wing discs described in E-G. Error bars indicate mean±s.d., and significance of differences for different developmental stages of *nub-GAL4:UAS-Dcr2/+* is relative to its 72h stage and indicated as ns in blue, the significance of differences relative to *nub-GAL4:UAS-Dcr2/+* at same developmental stage, calculated by t tests, is indicated by ns or black asterisks. **(I-K)** Third instar larval wing discs showing labeled clones using Raeppli technique from control (H, n=13 discs), *RNAi*-rok (I, n= 14 discs) *RNAi*-ft (J, n=19 discs). Scale bar=50 µm. **(L)** Quantification of clone roundness from discs described in I-K, by violin plot, N=132 (control), 122(RNAi-rok), and 243(RNAi-fat) clones. Error bar indicates mean±s.d., the significance of differences relative to control, calculated by one-way ANOVA, is indicated by asterisks.

During normal development, growth within the wing pouch is preferentially oriented in a proximal-distal direction, as revealed by the elongation of marked clones ^40^. *ds* or *fat* mutants randomize the orientation or growth, resulting in rounder clones that are no longer oriented along the proximal-distal axis ^40,56^. To evaluate the contribution of cytoskeletal tension to oriented growth, we examined clones of cells in *nub-Gal4 UAS-RNAi-rok* wing imaginal discs and compared them to clones in wild-type and in *nub-Gal4 UAS-RNAi-fat* wing discs. Clones were marked using the Raeppli technique, which generates clones labelled with one of four different fluorescent proteins ^57^. Clones of cells in *rok* knock down discs were rounder than those in wild-type discs, supporting a key role for cytoskeletal tension in orienting growth during wing development (Figure 6I, J, L). However, they were not as round as those in fat RNAi discs (Figure 6K, L), consistent with the observation that *rok* RNAi wings are not as round as *fat* RNAi wings.

### Increased cytoskeletal tension modulates the shape of *dachsous* knock-down wings

Observations that loss of *ds* alters stress patterns in wing discs, together with observations that direct manipulation of cytoskeletal tension alters clone growth and wing shape, led us to investigate whether manipulating cytoskeletal tension could modify the consequences of *ds* knockdown on wing shape. We found that knocking down *rok* together with *ds* did not further increase wing roundness (Figure 7A-D,G). Conversely, increasing Rok activity by expressing Rok.CA in *ds* knockdown wings partially rescued wing shape, as wings were elongated compared to *ds* knockdown wings (Figure 7C, E-G). This implies that the reduced cytoskeletal tension in *ds* knockdown wings contributes to the effect of *ds* on wing shape.

**Figure 7:**
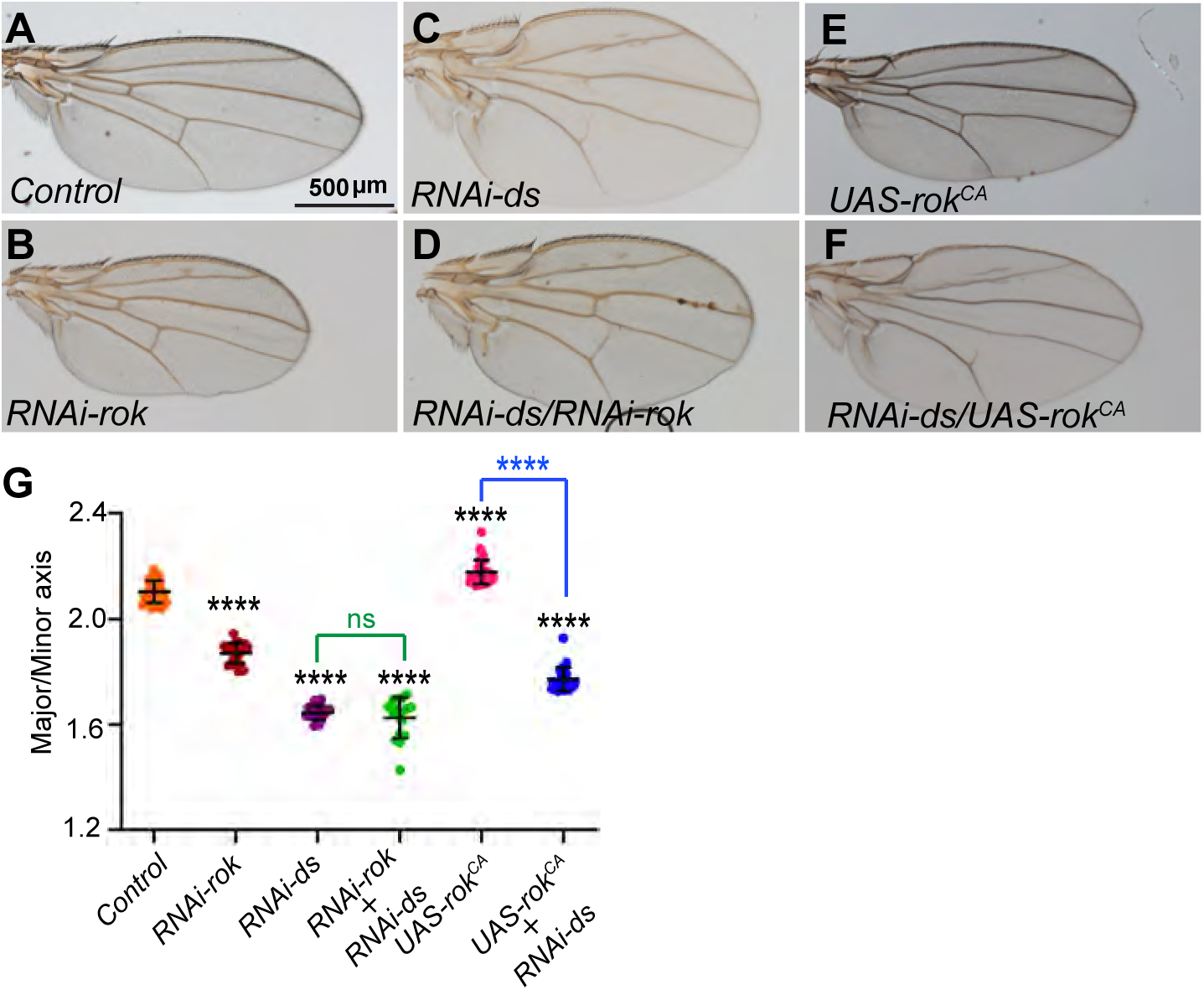
Increasing cytoskeletal tension partially rescues *ds* wing shape. **(A-F)** Male wings from *nub-GAL4:UAS-Dcr2/+* control (A; n=32), *RNAi-rok* (B; n=23), *RNAi-ds* (C; n=19), *RNAi-rok, RNAi-ds* (D; n=17), *UAS-rok^CA^* (E; n=33), and *UAS-rok^CA^, RNAi-ds* (F; n=27). Scale bar=500 µm. **(G)** Histogram quantifying shape for adult wings described in A-F. Error bar indicates mean±s.d., the significance of differences compared to control by one-way ANOVA is indicated by black asterisks and between *RNAi-rok* and *UAS-rok^CA^* on *ds* knock-down is shown in green and blue, respectively.

We also compared the impact of reduced cytoskeletal tension on wings from animals with impaired Hippo signaling by expressing *rok* RNAi in *ex* mutant or *hpo* knockdown flies. This further increased the roundness of the adult wing (Figure S5A-E). Moreover, decreasing Rok levels in *ex* mutants increased the D-V/A-P ratio in wing discs throughout larval development (Figure S5F,G). These observations further support the inference that Ds-Fat signaling impact wing shape through effects on cytoskeletal tension that are distinct from its connection to Hippo signaling.

## Discussion

Ds-Fat signaling controls organ shape from *Drosophila* to mammals, but the mechanisms involved have remained poorly understood ^17–19^. One of the best studied examples of this is the *Drosophila* wing, which normally has an elongated shape, but becomes more rounded upon mutation or knockdown of genes involved in Ds-Fat signaling ^4–6,8^. Pioneering studies suggested that this was mediated through controlling the orientation of mitotic spindles to orient growth along the proximal-distal axis ^40,41^. However, the observation that randomizing spindle orientation did not alter wing shape or growth orientation within the wing disc argued against this ^42^. Our observation that stress patterns and myosin distribution are altered in *ds* mutant wing discs, together with observations that direct manipulation of cytoskeletal tension modulates wing shape, suggest that Ds-Ft signaling modulates wing shape primarily through regulating tissue tension during larval growth. Elongation of the wing primordia along the proximal distal axis, which has been revealed by cellular dynamics to include a combination of oriented cell rearrangement, cell shape changes, and cell divisions, is believed to occur in response to normal stress patterns ^43^. The effect of Ds-Fat signaling on spindle orientation could thus be understood as an indirect consequence of the altered stress patterns that occur when genes in this pathway are inactivated.

Our results also emphasize that while the wing undergoes dramatic morphogenesis that reshapes it from imaginal disc to wing during pupal development, and Ds and Fat are expressed and actively influence PCP during pupal development ^45–47^, their influence on wing shape occurs prior to this, during the larval growth phase. This was confirmed functionally, by temperature shift experiments establishing that *ds* and *fat* are required during larval development to regulate wing shape. In addition, we observed that their effect on the shape of the wing pouch in the larval imaginal disc prefigures their effect on the shape of the adult wing. That is, despite the dramatic reshaping of wing tissue during metamorphosis, the altered shape of the future wing is evident in the larval disc in the relative increase in length of DV boundary (wing circumference) as compared to the length of AP boundary (wing length). This observation is consistent with the idea that adult wing shape is preprogrammed in larval wing disc morphology ^58^.

Ds-Fat signaling modulates distinct, Dachs-dependent, downstream processes to regulate growth, PCP and morphogenesis. We found that Ds-Fat alters the shape of the developing wing pouch from the earliest time that it becomes visibly outlined by Wg expression, at early third instar. This effect of Ds-Fat was shared by inactivation of genes that specifically impair Hippo signaling. However, the initial effect of Hippo pathway genes on wing disc shape was lost during disc growth, and the adult wings that formed from discs where these genes were impaired are not as round as *ds* or *fat* mutant wings. Thus, we conclude that effects on Hpo signaling are not able to explain the effect of Ds-Fat on wing shape. This conclusion is further supported by the observation that wing shape in animals with compromised Hippo signaling in the wing remains sensitive to decreases in cytoskeletal tension generated by *rok* RNAi, whereas *ds* wing shape was not further altered by *rok* RNAi. Ds-Ft signaling also connects to canonical PCP, through physical interaction of Ds and Dachs with the Sple isoform of Pk-Sple ^32,33^. However, as mutation or knock down of core PCP components, including *pk* or *sple*, does not affect wing shape ^59^, this cannot account for the effect of Ds-Fat on wing shape. Finally, Ds-Fat have also been proposed to regulate tissue tension through their effects on localization of Dachs ^36^, which is a myosin family protein that can bind F-actin ^5,60^. The potential ability of Dachs to affect junctional tension has previously been suggested to contribute to effects of Ds-Fat signaling on the shape of the notum during pupal morphogenesis ^36^, and we suggest that effects of Dachs on tissue tension are also likely to explain the influence of Ds-Fat on wing shape. In the notum, however, Dachs was inferred to act directly during pupal morphogenesis, in contrast to our observations that Ds-Fat makes its key contributions to morphogenesis during larval development.

As Dachs localization is polarized in the developing wing imaginal disc ^5^, the influence of Ds-Fat signaling may in part depend upon this polarization. Consistent with this, junctional tension in much of the wing is normally higher along circumferential junctions (where Dachs accumulates) than along radial junctions ^61,62^, and we observed a Ds-dependent stress anisotropy in the proximal wing. However, we also see global effects on the overall distribution of tissue tension contributing to shaping the developing wing. This is emphasized first by the broad changes in myosin distribution and tissue tension revealed by imaging and laser cutting. The importance of tissue wide effects on tension is also emphasized by the non-autonomous nature of the effects of *dachs* on growth orientation. Thus, while marked clones of cells within *dachs* mutant discs lack normal growth orientation, clones of cells mutant for *dachs* in otherwise wild-type discs exhibit normal growth orientation ^5,41^. The observation that wings can be made rounder by decreasing cytoskeletal tension, and more elongated by increasing cytoskeletal tension, also argue for an important contribution of global tension patterns to wing shape. Nonetheless, the observation that influence of *ds* on wing shape could only be partially rescued by a broad increase in tension suggests that the normal patterns and polarization of tension within the developing wing disc is also important form normal shape. Finally, we emphasize that the observation that Ds-Fat signaling controls wing morphogenesis through regulation of tissue tension may also apply to other contexts where Ds-Fat signaling controls morphogenesis.

## MATERIALS AND METHODS

### Drosophila genetics

All flies were kept on standard cornmeal fly food supplemented with yeast and agar. Stocks were maintained at room temperature. Fly crosses were performed at 25°C unless otherwise specified. The stocks that were used in this study include: *w*^1118^(control)*, ds^UA^*^071^^63^*, ds^36D^* ^64^*, RNAi-ds* (vdrc 36219), GS-ds, *d^GC^*^13^ and *d*^210^ ^5^, *ft^G-rv^*(BDSC#1894), *ft*^8^ (BDSC#44257), *RNAi-ft* (vdrc#9396), UAS-fat ^65^, *ex^e^*^1^ (BDSC#44249)*, RNAi-hpo* (vdrc 104169)*, wts^P^*^2^ ^66^*, wts^X^*^1^ ^67^, UAS-wts ^55^, Ecad:GFP ^68^ ^36^, sqh-sqh:GFP ^69^*, nub-Gal4* ^70^ ^55^*, hh-Gal4* ^44^, tub-Gal80^ts^ (BDSC#7017), UAS-dcr2 ^71^, *RNAi-rok* (vdrc104675), UAS-rok^CA^ (BDSC#6668), Raeppli-CAAX[67E] (BDSC#55083), UAS-InR^act^ (BDSC#8263), UAS-mCD8:RFP (gift of G. Morata, Universidad Autónoma de Madrid, Madrid). *inv-BFP ap-BFP* were created by cloning enhancer sequences driving wing imaginal disc expression of *inv* (R88c04) and *ap* (R42A06) ^72^ upstream of 2xmTagBFP in place of Gal4 in pBPGUw (addgene 17575) and isolating third chromosome insertions.

### Temperature Shift experiments

For temporal knockdown experiments, we used the *nub-GAL4* driver line in combination with *tub-Gal80^ts^* crossed to *RNAi-ft* (vdrc 9396) or *RNAi-ds* (*vdrc 36219*). Crosses and progeny were kept at either 18 °C or 29 °C, following the temperature shift scheme described in Figure 1.

### Adult wing imaging and analysis

Adult male wings were dissected in isopropanol and mounted in 4:1 Canada Balsam:Methyl Salicylate and imaged using a Zeiss Axioplan2 microscope and a Progress camera. The major and minor axes, as well as the area of adult wings, were measured by manually tracing digital wing images by using ‘Free hand selection tool’ in Fiji (Schindelin et al., 2012) with ‘Fit Ellipse’ ‘Area’ and ‘Shape Descriptors’ options selected in ‘Set measurement tool’ selections.

### Pupal wing Imaging

For pupal wing imaging, larvae of the desired genotype were scored at the late 3^rd^ instar stage and kept at 25°C, scored as 0h After puparium formation (APF) when they formed white prepupae and imaged at 6h, 12h and 18h APF. Prior to imaging at each stage, pupae were gently removed from vials using a moistened brush (Princeton Velvetouch Series 3950 Synthetic Round Brush, Size 1) and aligned on a glass slide with one pupal wing oriented toward the objective. Pupae were aligned using a stereo microscope (Zeiss Stemi SV11 Apo) equipped with fluorescence illumination (Excelitas X-Cite Series 120Q). Once aligned, the pupae adhered to the slide within a few seconds and were ready for imaging. Pupal wings were imaged through the pupal case using a Leica TCS SP8 Confocal Microscope.

### Immunostaining and Microscopy

To obtain wing discs at different stages, the flies were transferred to new vials for 4 to 6h, and larvae were dissected at 72h, 84h, 96h, 108h and 120h After Egg Laying (AEL). For most experiments, wing discs were fixed for 15 min in 4% paraformaldehyde at room temperature (RT), whereas Sqh:GFP discs were fixed for 12 min in 4% paraformaldehyde at RT. Fixed larval tissues were rinsed twice with PBT [1xPBS with 0.1% (v/v) Triton-x-100 + 1% (w/v) BSA + 0.01% (v/v) Na-azide], then washed 3 times 15 minutes each in PBT at RT, incubated for 30 min in blocking solution [PBT + 5% (v/v) Donkey serum], and incubated with primary antibodies overnight at 4 °C with gentle mixing. Primary antibodies used were rat anti-E-cadherin (Developmental Studies Hybridoma Bank, DCAD2-c; 1:200), mouse anti-Wg (Developmental Studies Hybridoma Bank, 9A4-c; 1:300), rabbit anti-Dcr2 (Abcam, ab4732; 1:1000). Sqh localization was monitored using sqh:GFP (Royou et al., 2004). Discs were then rinsed twice with PBT, then washed 4 times 15 minutes each in PBT, incubated for 30 min in blocking solution, and incubated with secondary antibodies for 2 hours at RT with gentle mixing, with tubes wrapped in aluminum foil. Secondary antibodies were from Jackson ImmunoResearch Laboratories, Invitrogen and Biotium (20137). Discs were then rinsed twice with PBT, washed 2 times 15 minutes in PBT, incubated 15 min with Hoechst 33342 (Invitrogen, H3570) to label DNA, rinsed twice with PBT, then washed 2 times 15 minutes each in PBT. Wing imaginal disc were dissected using fine forceps, and mounted onto a microscope slide in Vectashield (Vector Laboratories, H-1000). Images were collected on the Leica SP8 confocal microscope.

### Wing pouch shape analysis

To calculate the dorsal-ventral axis (D-V)-anterior-posterior axis (A-P) ratio for wing pouch shape analysis, the wing discs were stained with mouse anti-Wg, and the wing pouch was defined inside the inner ring of Wg expression. The D-V boundary and an approximate A-P axis was manually traced inside the inner ring of Wg expression using the ’Freehand Line’ tool in Fiji. The aspect ratio of the wing pouch was calculated by dividing the length of the D–V axis by the length of the A–P axis.

### Raeppli Clones

To generate Raeppli clones, *nub-Gal4 UAS-dcr2> Raeppli-CAAX-67E* flies were crossed with either *w*^1118^ (control), *RNAi-rok,* or *RNAi-ft* and then maintained at 25°C. Larvae were heat-shocked in a water bath at 36 °C for 8 minutes between 60–72 hours AEL, then transferred to 29°C. Larvae were dissected 36 hours after heat shock to analyze Raeppli clones. To quantify Raeppli clone shape, clone boundaries were manually traced and analyzed using the Shape Descriptors function in Fiji/ImageJ.

### Statistical Analysis

Statistical significance was examined using GraphPad Prism software by performing Student’s *t*-test for comparisons between two groups or one-way ANOVA for multiple groups, with *p* < 0.05 considered statistically significant. All quantifications are presented as the mean ± SD. For all statistical tests, ns indicates P>0.05, * indicates P≤0.05, ** indicates P≤0.01, *** indicates P≤0.001, **** indicates P≤0.0001.

### Image Analysis

For the Sqh:GFP fluorescence intensity heat map, confocal image stacks were processed using a custom MATLAB script (Alégot et al., 2019; Pan et al., 2018) to project the apical surface onto a 2D plane, based on the maximal brightness of E-Cadherin. The intensity of the channel of interest (Sqh:GFP) is normalized over the reference channel (E-cad) intensity and represented by the heat map. To quantify cell shapes, confocal image stacks were processed using a custom MATLAB script (Alégot et al., 2019; Pan et al., 2018) to generate flattened projections. The Tissue Analyzer plugin in Fiji was then used to segment individual cells based on the E-cad signal. Cell eccentricity was calculated using the Quantify Polarity software ^73^.

### Live imaging and Circular Ablation

For circular ablation experiments, wing discs at 108h were dissected and cultured in live imaging media by following the protocol established by Dye et al., 2017 ^43^. This uses Grace′s Insect Medium (With L-glutamine, without sodium bicarbonate) (Sigma-Aldrich, G9771) dissolved in water with pH adjusted to 6.6–6.7 and filter sterilized (Thermo Scientific # 450-0020). Prior to each experiment, the Grace′s Insect Medium stock solution was supplemented with 5% fetal bovine serum (Thermo Fisher, 10082147), 100× penicillin-streptomycin (Thermo Fisher, #15070063), and 10 nM 20-hydroxy-ecdysone (Sigma, H5142). Wing disc dissection and mounting was done as described previously ^42^;^74^. Circular ablation was performed using an Andor Dragonfly500 Spinning Disk Confocal with a MicroPoint ablation system. Circles of 10 μm radius were cut using Micropoint settings of 60% Laser power, Repetition rate 16Hz, and 2 Repeats. Roundness and maximum expansion area were measured 1 minute after ablation using Fiji/ImageJ. Each ablation was performed on a separate wing disc. Graphs were made using GraphPad Prism, and images were arranged in Adobe Illustrator.

## Acknowledgments

We thank Joe Terzian and Nazir Qureshi for experimental assistance and analysis, and Shuguo Sun for creating inv-BFP ap-BFP flies. This research used antibodies from the Developmental Studies Hybridoma Bank, fly stocks from the Bloomington Drosophila Stock Center and Vienna Drosophila Resource Center, information from FlyBase, and microscopy resources of the Waksman Institute Shared Imaging Facility. This research was supported by National Institutes of Health grant GM131748 (KDI).

## Supplemental Figures

**Figure S1:**
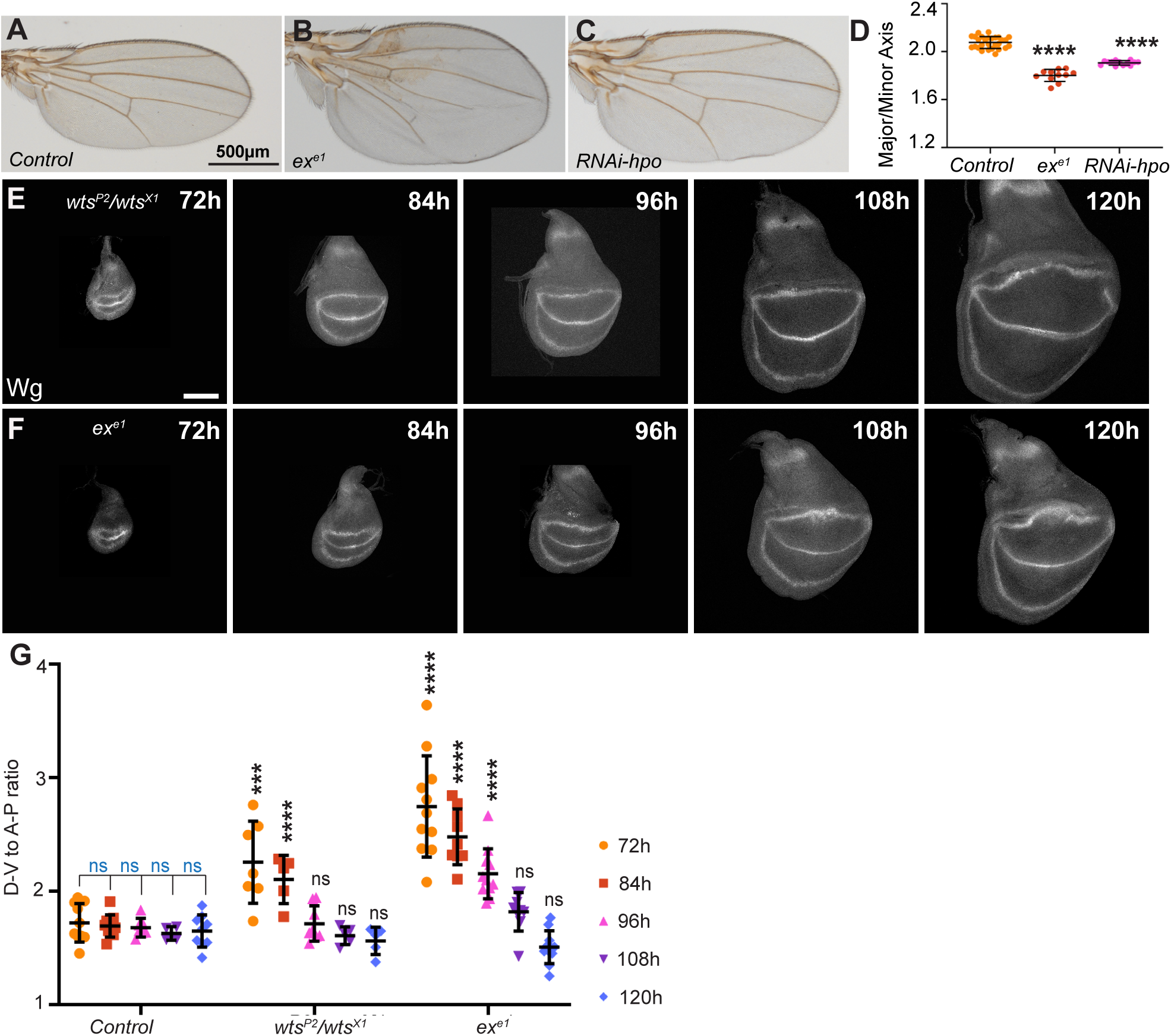
Influence of Hippo signaling on shape of larval wing pouch and adult wing. **(A-C)** Male wings from *w*^1118^ (A; n=35), *ex^e^*^1^*/ ex^e^*^1^ (B; n=11) and *nub-GAL4:UAS-Dcr2/RNAi-hpo* (C; n=13). Scale bar=500 µm. **(D)** Histogram quantifying shape for adult wings described in A-C. Error bar indicates mean±s.d., the significance of differences relative to *w*^1118^, calculated by one-way ANOVA, is indicated by asterisks. **(E, F)** Wing discs stained for Wg from **(E)** *wts^P^*^2^ /*wts^X1^*at 72h AEL (n=7), 84h AEL (n=5), 96h AEL (n=8), 108h AEL (n=5), and 120h AEL (n=5) **(F)** *ex^e^*^1^*/ ex^e^*^1^ at 72h AEL (n=11), 84h AEL (n=9), 96h AEL (n=11), 108h AEL (n=9) and 120h AEL (n=11). Scale bar=100 µm. **(G)** Histogram quantifying wing pouch shape for discs described in E,F. Error bar indicates mean±s.d., the significance of differences relative to *w*^1118^ at same developmental stage, calculated using one-way ANOVA, is indicated by asterisks or ns. Control data is the same as of shown in Figure 3A, D and used here for comparison.

**Figure S2:**
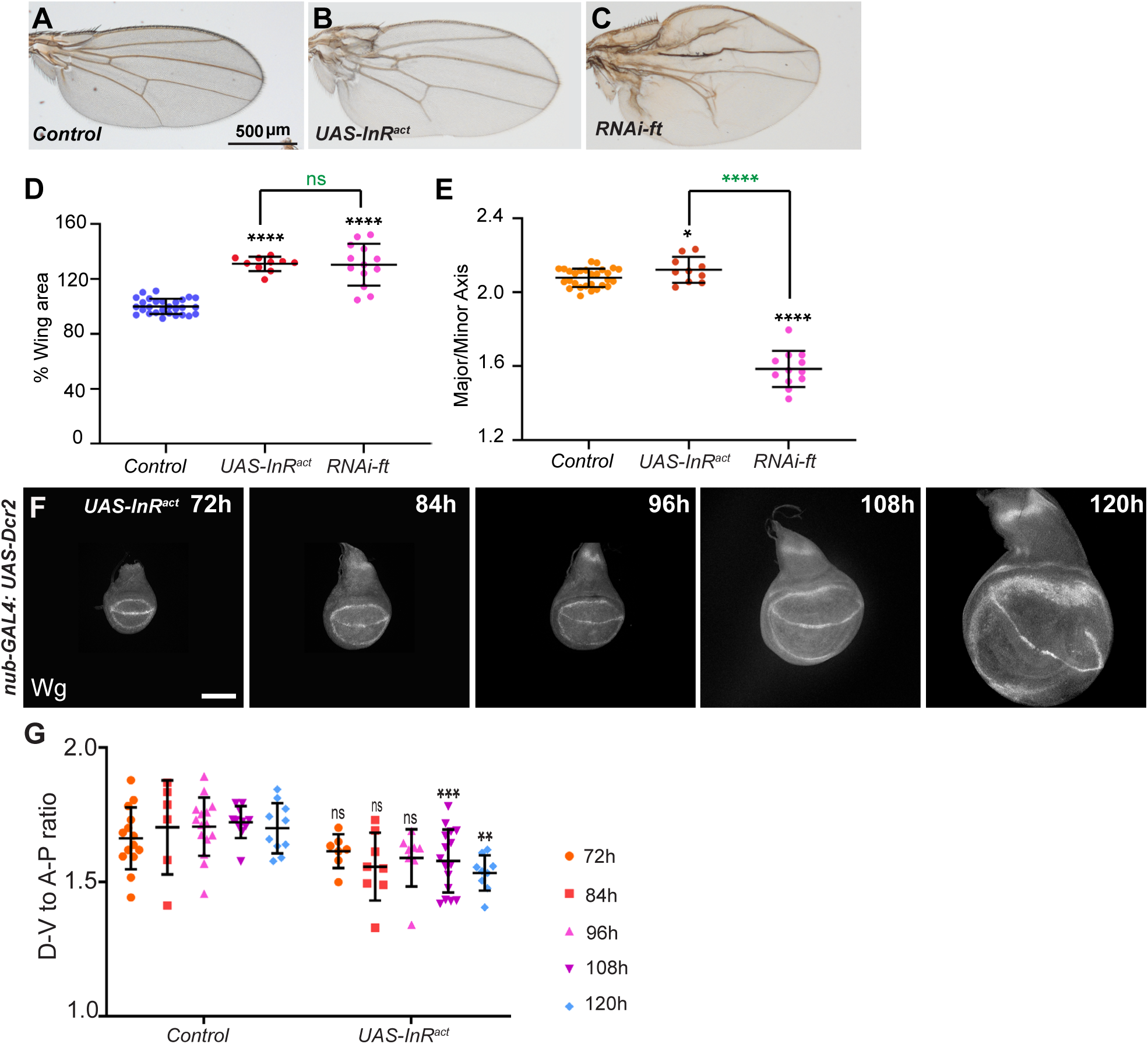
Influence of InR activation on wing shape. **(A-B)** Male wings from *nub-GAL4:UAS-Dcr2/+* as control (A; n=28), *nub-GAL4:UAS-Dcr2/UAS-InR^act^* (B; n=10), *nub-GAL4:UAS-Dcr2/RNAi-ft* (C; n=12). Scale bar=500 µm. **(D, E)** Histogram quantifying (D) wing area (E) wing shape, compared to *nub-GAL4:UAS-Dcr2/+* .Error bar indicates mean±s.d., the significance of differences relative to *nub-GAL4:UAS-Dcr2/+* is indicated by black asterisks and was calculated using one-way ANOVA on measurements from the number of adult wings indicated above and the significance of difference between *UAS-InR^act^* and *RNAi-ft* is indicated by ns or asterisks in green. **(F)** Wing discs stained for Wg from *nub-GAL4:UAS-Dcr2/UAS-InR^act^* at 72h AEL (n=7), 84h AEL (n=8), 96h AEL (n=8), 108h AEL (n=16), and 120h AEL (n=9). Scale bar=100 µm. **(G)** Histogram quantifying wing pouch shape. Error bar indicates mean±s.d., the significance of differences relative to *nub-GAL4:UAS-Dcr2/+* from the same developmental stage is indicated by ns/black asterisks and was calculated using t-tests on measurements from the number of samples indicated above. Control data is the same as of shown in Figure 6E, H and used here for comparison.

**Figure S3:**
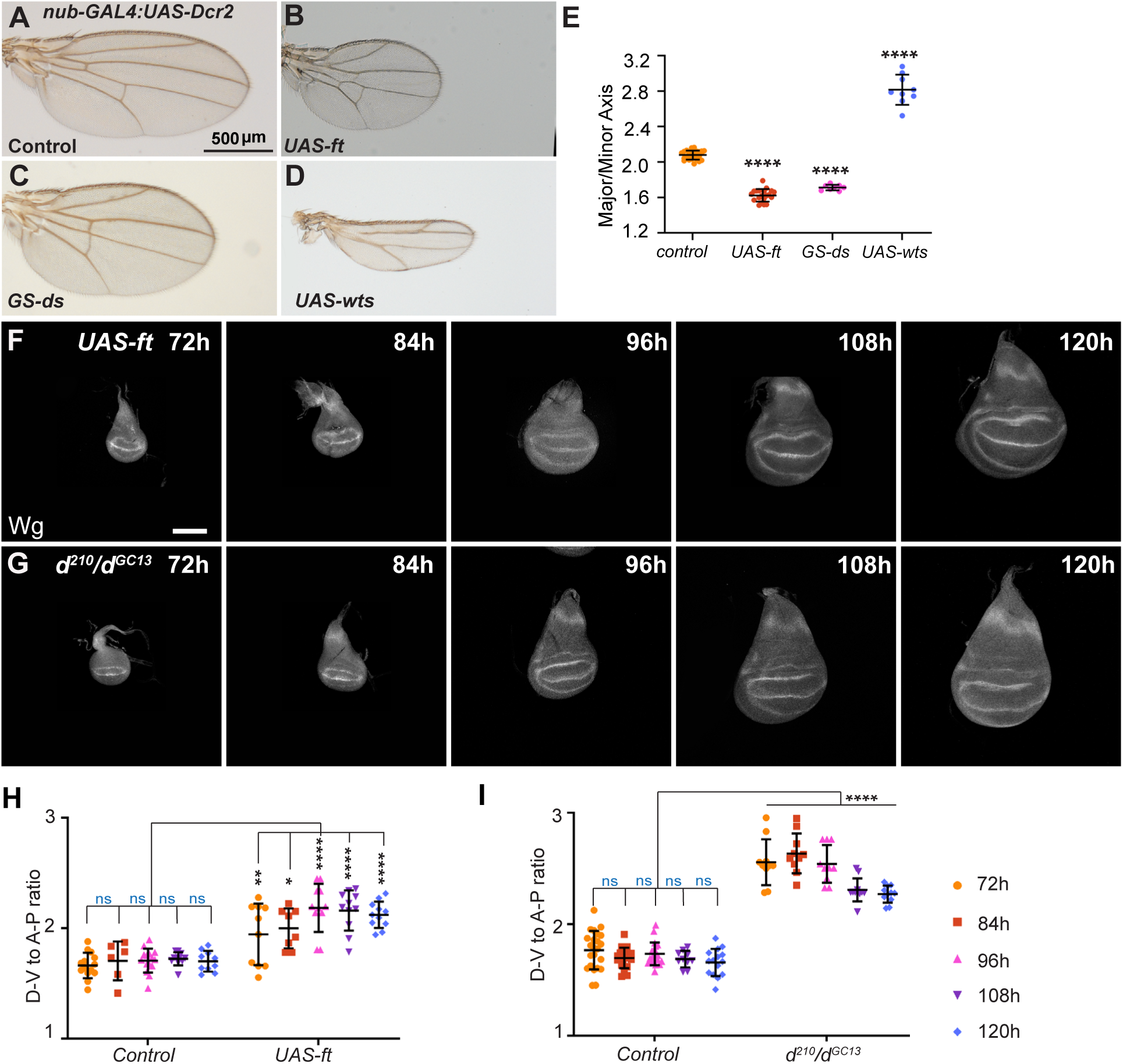
Influence of Fat overexpression or *dachs* mutant on wing pouch shape. **(A-D)** Male wings from *nub-GAL4:UAS-Dcr2/+* (A; n=28), *nub-GAL4:UAS-Dcr2/UAS-ft* (B; n=19), *nub-GAL4:UAS-Dcr2/GS-ds* (C; n=8), and *nub-GAL4:UAS-Dcr2/UAS-wts* (D; n=9) . Scale bar=500 µm. **(E)** Histogram quantifying wing shape for wings described in A-D. Error bar indicates mean±s.d., the significance of differences relative to *nub-GAL4:UAS-Dcr2/+*,calculated by one-way ANOVA, is indicated by asterisks. **(F, G)** Wing discs stained for Wg **(F)** from *nub-GAL4:UAS-Dcr2/UAS-ft* at 72h AEL (n=10), 84h AEL (n=8), 96h AEL (n=11), 108h AEL (n=11), and 120h AEL (n=11) **(G)** *d*^210^*/ d^GC^*^13^ at 72h AEL (n=10), 84h AEL (n=10), 96h AEL (n=10), 108h AEL (n=11), and 120h AEL (n=11). Scale bar=100 µm. **(H,I)** Histograms quantifying wing pouch shape for F and G, respectively. Error bars indicate mean±s.d., the significance of differences relative to *nub-GAL4:UAS-Dcr2/+* (control for H, same as shown in Figure 6E, H) and *w*^1118^ (control for I, same as shown in Figure 3A, D).

**Figure S4:**
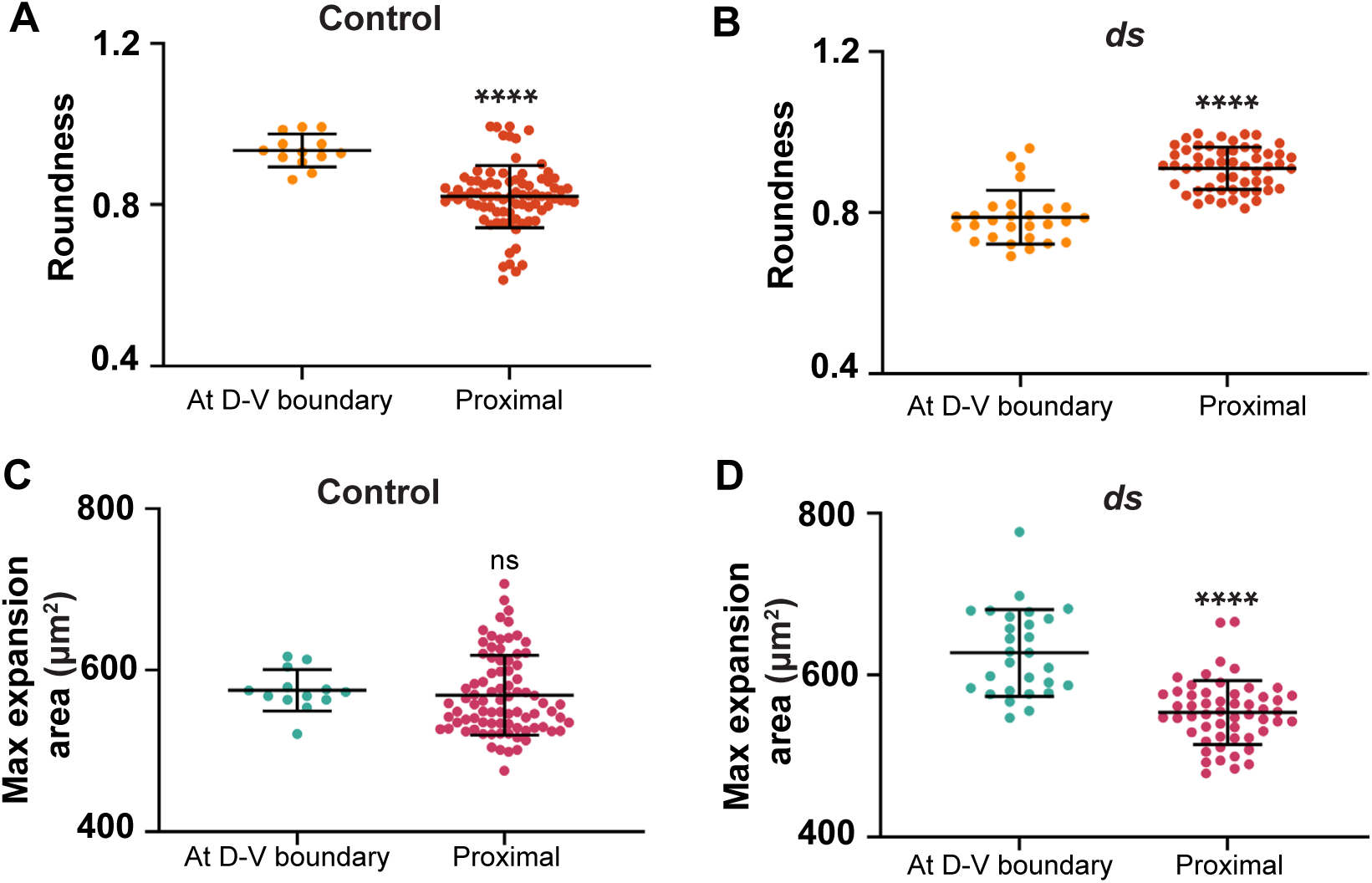
Comparison of the pattern of stress in control and *ds* mutants in different parts of the wing disc by circular ablation. **(A-D)** Histograms, based on data described in Figure 4, compare the shape (A, B) and size (C, D) of cut regions at their maximal expansion. Error bar indicates mean±s.d., the significance of differences is indicated by black asterisks and were calculated using t-tests.

**Figure S5:**
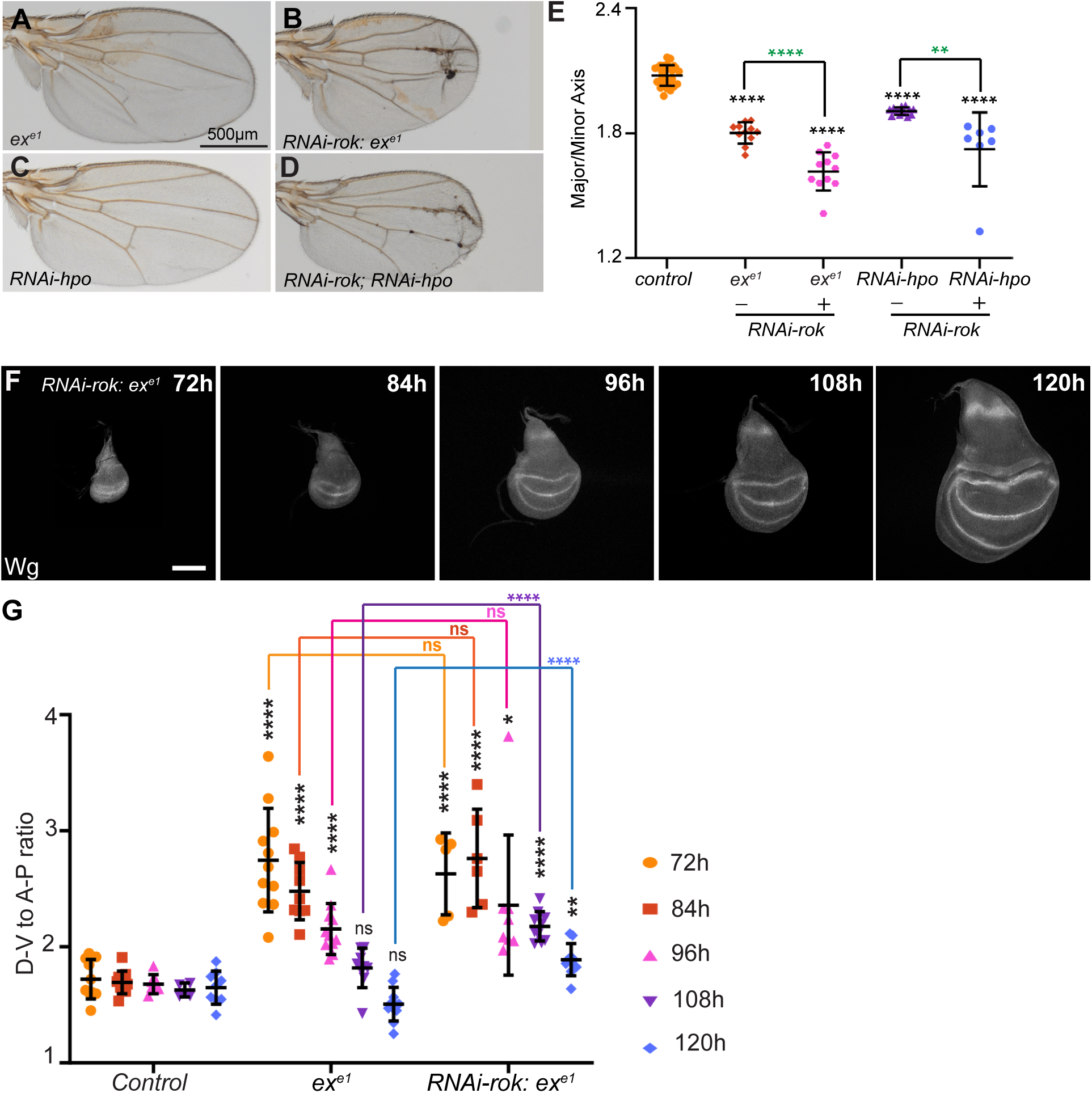
Decreased cytoskeletal tension modifies wing shape in Hippo pathway mutants. **(A-D)** Male wings from *ex^e^*^1^ (A; n=11), *RNAi-rok: ex^e^*^1^ (B; n=11), *RNAi-hpo* (C; n=13), and *RNAi-rok, RNAi-hpo* (D; n=7). Scale bar=500 µm. **(E)** Histogram quantifying shape for wings described in A-D. Error bar indicates mean±s.d., the significance of differences relative to control, calculated by one-way ANOVA, is indicated by black asterisks, and differences between presence or absence of *RNAi-rok* is indicated by green asterisks. **(F)** Wing discs stained for Wg from *RNAi-rok: ex^e^*^1^ at 72h AEL (n=5), 84h AEL (n=6), 96h AEL (n=8), 108h AEL (n=10), and 120h AEL (n=8). Scale bar=100 µm. **(G)** Histogram quantifying wing pouch shape for (F). Error bar indicates mean±s.d., the significance of differences relative to control (control same as shown in Figure 3A, D; *ex^e1^*same as shown in Figure S1F, G) is indicated by ns/black asterisks and was calculated using t-tests on measurements from the number of samples indicated above. Comparison between the with and without RNAi-rok group was done by a t-test, and the significance of differences is presented between the groups as color coded.

